# Speech intelligibility changes the temporal evolution of neural speech tracking

**DOI:** 10.1101/2022.06.26.497639

**Authors:** Ya-Ping Chen, Fabian Schmidt, Anne Keitel, Sebastian Rösch, Anne Hauswald, Nathan Weisz

**Affiliations:** Centre for Cognitive Neuroscience, University of Salzburg, 5020 Salzburg, Austria; Department of Psychology, University of Salzburg, 5020 Salzburg, Austria; Psychology, School of Social Sciences, University of Dundee, DD1 4HN Dundee, UK; Department of Otorhinolaryngology, Paracelsus Medical University, 5020 Salzburg, Austria; Neuroscience Institute, Christian Doppler University Hospital, Paracelsus Medical University, 5020 Salzburg, Austria

**Keywords:** vocoded speech, temporal response function, coherence, FOOOF, MEG

## Abstract

Listening to speech with poor signal quality is challenging. Neural speech tracking of degraded speech has been used to advance the understanding of how brain processes and speech intelligibility are interrelated, however the temporal dynamics of neural speech tracking are not clear. In the present MEG study, we thereby exploited temporal response functions (TRFs) and generated signal-degraded speech to depict the temporal evolution of speech intelligibility modulation on neural speech tracking. In addition, we inter-related facets of neural speech tracking (e.g., speech envelope reconstruction, speech-brain coherence, and components of broadband coherence spectra) to endorse our findings in TRFs. Our TRF analysis yielded marked temporally differential effects of vocoding: reduction of intelligibility went along with large increases of early peak responses (∼50-110 ms, M50_TRF_), but strongly reduced responses around 175-230 ms (M200_TRF_). For the late responses 315-380 ms (M350_TRF_), the maximum response occurred for degraded speech that was still comprehensible then declined with reduced intelligibility. Furthermore, we related the TRF components to our other neural “tracking“ measures and found that M50_TRF_ and M200_TRF_ play a differential role in the shifting center frequency of the broadband coherence spectra. Overall, our study highlights the importance of time-resolved computation and parametrization of coherence spectra on neural speech tracking and provides a better understanding of degraded speech processing.

**Highlights:**

- We use MEG to show that speech intelligibility differentially impacts the temporal evolution of neural speech tracking.
- TRF responses around 200 ms show the strongest relationship with behaviour.
- Relating TRF effects to parameterized coherence spectra using FOOOF suggests that M50_TRF_ and M200_TRF_ reflect shifts in which speech features are tracked over time.

## 1. Introduction

Listeners process speech under a variety of adverse conditions in daily life (Mattys et al., 2012), which could be external (e.g., “cocktail party“) or internal (e.g., hearing damage). However, how the neural dynamics of speech processing in such challenging conditions evolve remains unclear. A popular approach to systematically manipulate speech intelligibility–and possibly mimicking the experience of cochlear implant (CI) users–in normal hearing individuals is by using noise-vocoded speech (Friesen et al., 2001; Rosen et al., 2013). This approach filters the speech signal into a given number of channels or bands (typically between 1 and 16 channels), resulting in reduced spectral information while temporal information is preserved. This manipulation allows to parametrically modulate speech intelligibility (Shannon et al., 1995) and to relate features of the signal to neural activity.

The high temporal resolution of electroencephalography (EEG) and magnetoencephalography (MEG) can be exploited to quantify how temporal fluctuations of speech and neural signals align together, a process often described as “neural speech tracking“. Most studies (e.g., Ding & Simon, 2012; Fiedler et al., 2019; Zion Golumbic et al., 2013), including those investigating the effects of vocoding, have focused on the speech envelope. Since the envelope of e.g. a target speaker can be easily obtained, neural tracking using M/EEG is an attractive option to study brain activity also during complex listening situations such as background noise or multi-speaker scenarios. Speech-brain coherence is likely the most established measure of neural tracking (Hauswald et al., 2020; Peelle et al., 2013; Schmidt et al., 2021). However, system identification approaches, such as temporal response functions (TRFs; Crosse et al., 2016; Ding et al., 2014; Kraus et al., 2021) and stimulus reconstruction (Nogueira et al., 2019; Verschueren et al., 2019), have been gaining a lot of popularity. TRFs especially yield interesting temporal information on the relationship between stimulus and neural activity, which goes beyond the “static” coherence measure. Interestingly, these measures are rarely reported together.

Speech envelope reconstruction yielded higher accuracies for healthy listeners as compared to CI users and was also increased for attended speech (Nogueira et al., 2019). Additionally, speech envelope reconstruction works better in CI individuals with good speech rehabilitation (Nogueira et al., 2019; Verschueren et al., 2019). Overall, these findings suggest stronger neural tracking relates to higher speech intelligibility. However, when studying individuals with hearing loss, Decruy et al. (2020) observed higher envelope reconstruction accuracy in participants with mild to severe sensorineural hearing loss than age-matched normal hearing adults. Studying TRFs during vocoded speech has shown stronger responses at early latencies (∼50 ms) for less intelligible speech (Ding et al., 2014; Kraus et al., 2021) and at later latencies for attended speech (Kraus et al., 2021). The notion that the mapping of speech intelligibility onto neural tracking is not straight-forward is also supported by studies using speech-brain coherence. For example, Peelle et al. (2013) and Schmidt et al. (2021) showed that the coherence decreased linearly as the speech intelligibility declined, whereas Hauswald et al. (2020) showed an inverted-U-shaped pattern. In the latter study, the highest coherence was observed in the still comprehensible, degraded speech rather than the clear speech. Computing cross-correlation between speech envelopes and neural activities, Millman et al. (2015) and Baltzell et al. (2017) did not observe an effect of speech intelligibility, whereas an effect of prior knowledge was found in Baltzell et al. (2017). Overall, current findings using different “flavors“ of neural tracking do not point to a consistent effect of vocoded speech.

Here, we utilized MEG and had our participants listen to speech at different vocoding levels. We applied the TRF method to explore how 6 levels of vocoded speech (Figure 1) modulate the neural speech tracking with fine-grained temporal resolution. As expected, participants’ behavioral performance declined as speech intelligibility deteriorated. Interestingly, our TRF results showed that degraded speech modulates the neural speech tracking at three intervals in differential ways: Responses at 50-110 ms (M50_TRF_) and 175-230 ms (M200_TRF_) generally captured whether speech was vocoded or not. Late responses around 315-380 ms (M350_TRF_) were strongest at medium vocoding levels as compared to clear and less intelligible speech. To complement our findings in TRF, we also computed speech envelope reconstruction, speech-brain coherence, and periodic components (peak center frequency, peak bandwidth, peak height) and aperiodic components (exponent, offset) of broadband coherence spectra (Schmidt et al., 2021). Relating the TRF components to those neural measures, we found that M50_TRF_ and M200_TRF_ were correlated with the center frequency of the broadband coherence spectra in a different direction. These results suggest that changes in TRF sensitive to speech intelligibility decreases reflect not only sensory gain or general attentional modulations but also shifts in which speech features are tracked.

**Figure 1.**
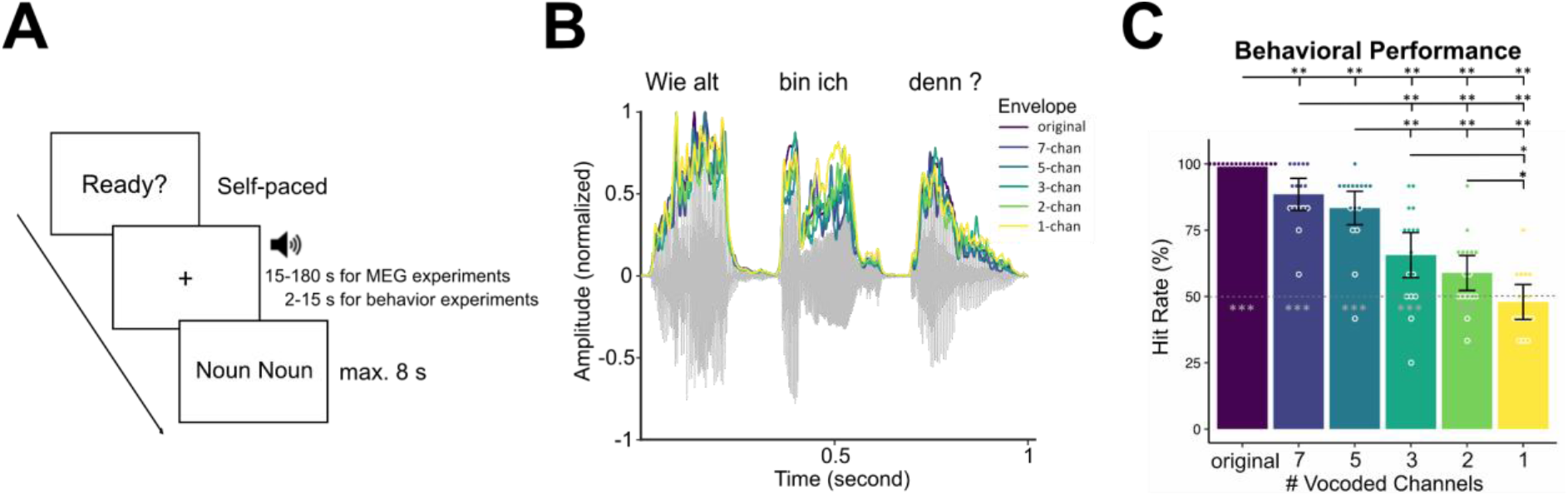
(**A**) Example trial of the MEG and behavioral experiment. Participants started self-paced and listened to the audiobook with eye fixation on the cross. At the end of the trial, two nouns were presented on the screen. Participants had to choose which noun was the one they heard from the last sentence. (**B**) An exemplary original audio segment with the corresponding envelopes and with the envelopes from the other vocoded conditions. The envelope from the original audio is essentially identical to the envelopes from the vocoded audio on power level.(**C**) The hit rate declined as the speech intelligibility decreased. Bars represent 95% confidence intervals. *p*_*fdr*_ < .05*, *p*_*fdr*_ < .01**, *p*_*fdr*_ < .001***

## 2. Materials and Methods

All data used in this study were also reported as Study 2 in Hauswald et al. (2020). The duration of the MEG experiment was ∼75 mins (preparation included) and the behavioral experiment was ∼7 mins. The behavioral experiment was also conducted in the MEG chamber without recording of MEG signals.

### 2.1 Participants

Twenty-four individuals [11 females, mean age = 26.4 ± 5.7 (SD), age range = 18-45 years] were recruited in the MEG experiment. They were native German speakers, reported normal hearing and no history of psychological or neurological disease, and were eligible for MEG recordings (i.e., without ferromagnetic metals in or close to their bodies). Sixteen of these individuals [7 females, mean age = 27.3 ± 6.8 (SD), age range = 18-45 years] also participated in the behavioral experiment. All the participants provided informed consent, and were compensated monetarily or with course credit. The recruitment and experiment procedure was in accordance with the Declaration of Helsinki and approved by the Ethics Committees of the Department of Psychology, University of Salzburg.

### 2.2 Stimuli

In both the MEG and behavioral experiments, participants were instructed to listen to audio files from audiobooks and to maintain visual fixation on a cross centered on screen for the duration of each trial (Figure 1A). We used a within-subjects design. There were six conditions in both experiments: original, 7-, 5-, 3-, 2-, or 1-channel (ch) noise-vocoded (Figure 1B). The speech vocoding was done using the vocoder toolbox (Gaudrain, 2016) for MATLAB. For the 7-, 5-, 3-, 2-, 1-ch vocoding levels, the waveform of each audio stimulus was passed through two Butterworth filters with a range of 200-7000 Hz and then further bandpass-filtered into the corresponding frequency analysis bands. For each band, a sinusoidal carrier was generated, and the frequency of the sine wave was equal to the center frequency of the analysis filter. Amplitude envelope extraction was done with half-wave rectification and low-pass filtered at 250 Hz. The amplitude-modulated noise bands from all the channels were then combined to produce the vocoded speech. The root mean square of the resulting signal was adjusted to that of the original signal.

In the MEG experiment, 24 audio files were created from recordings of a female German native speaker reading Goethe’s “Das Märchen” (1795). Lengths of stimuli varied between 15 s and 180 s, with two stimuli of 15 s, 30 s, 60 s, 120 s, 150 s, and 12 of 180 s. Each stimulus ended with a two-syllable noun within the last four words in the last sentence. The assignment of stimuli to conditions in the MEG experiment was controlled in order to obtain a similar overall length of stimulus presentation (∼400 s) in each condition. The order of the stimuli did not follow the order of the original story. The order of conditions was pseudorandomized according to a latin square design. Three stimuli were presented in one block after which a short break was offered, resulting in eight blocks. Each block was followed by a self-determined break. At the end of each auditory stimulus, participants were required to choose the noun in the last sentence from two two-syllable nouns on the screen. Following the response, they could self-initiate the next trial via a button press. The syllable rate of the 24 audio files varied between 4.1 and 4.5 Hz with a mean of 4.3 Hz, which was computed with a custom script from de Jong & Wempe (2009) using Praat (Boersma & Weenink, 2019).

In the behavioral experiment, 48 audio files were created from recordings of another female German native speaker reading a German version of Antoiné St. Exupery’s “The little prince” (1943) written by Grete and Josef Leitgeb (1956). Each stimulus contained one sentence (length between 2-15 s) and ended with a two-syllable noun within the last four words. Similar to the MEG experiment, participants were asked to choose the last noun they heard between two nouns on the screen. Again, the order of conditions was pseudorandom.

Stimulus presentation was controlled using a MATLAB-based objective psychophysics toolbox (Hartmann & Weisz, 2020) built based on the Psychtoolbox (Brainard, 1997; Kleiner et al., 2007; Pelli, 1997). Auditory stimuli were presented binaurally using MEG-compatible pneumatic in-ear headphones (SOUNDPixx, VPixx technologies, Canada). The trigger-sound delay of 16 ms was measured via the Black Box Toolkit v2 and was corrected during preprocessing of MEG data. Participants’ responses were acquired via a response pad (TOUCHPixx response box, VPixx technologies, Canada).

### 2.3 Data acquisition and analyses

#### 2.3.1 Extraction of the acoustic speech envelope

For computation of the temporal response functions and stimulus reconstruction, we extracted the acoustic speech envelope from all auditory stimuli using the Chimera toolbox (Smith et al., 2002), with which five frequency bands in the range of 200 to 4000 Hz were constructed as equidistant on the cochlear map. Sound stimuli were band-pass filtered (forward and reverse) in 5 bands using a 4th-order Butterworth filter. For each band, envelopes were calculated as absolute values of the Hilbert transform and were averaged across bands to obtain the full-band envelope. Envelopes for all 6 conditions were processed with this procedure and used for the following TRF, stimulus reconstruction, and coherence analysis.

To determine the envelope modulation rate, custom Matlab scripts from Ding et al. (2017) were used. The audio files were chunked into 10-s duration segments, resulting in 306 segments per condition. After calculating a Fast Fourier Transformation (FFT), the global maximum value of each 306 power spectrum was taken and then averaged as the envelope modulation rate. The envelope modulation rate of the 24 audio files in the original, 7-ch, 5-ch, 3-ch, 2-ch, and 1-ch vocoded condition was 5.61 Hz, 5.53 Hz, 5.27 Hz, 5.23 Hz, 4.93 Hz, and 4.83 Hz, respectively. A main effect of vocoding was observed [*𝒳*^2^(5) = 206.0, *p* = 1.83 × 10^−42^, Kendall’s W = 0.134, Friedman test], in line with Schmidt et al. (2021).

#### 2.3.2 MEG acquisition and preprocessing

MEG signals were recorded at a sampling rate of 1000 Hz using a 306-channel Triux MEG system (Elekta-Neuromag Ltd., Helsinki, Finland) with 102 magnetometers and 204 planar gradiometers in a magnetically shielded room (AK3B, Vakuumschmelze, Hanau, Germany). The MEG signal was online high-pass and low-pass filtered at 0.1 Hz and 330 Hz respectively. Prior to the recording, individual head shapes were digitized for each participant including fiducials (nasion, bilateral pre-auricular points) and at least 300 points on the scalp using a Polhemus Fastrak system (Polhemus, Vermont, USA). A signal space separation algorithm implemented in the Maxfilter software (version 2.215) provided by the MEG manufacturer was used to remove external noise from the MEG signal (mainly 16.6 Hz from Austrian local train power and 50 Hz plus harmonics from power line) and realign data across different blocks to an individual common head position (based on the measured head position at the beginning of each block).

Data analysis was done using the Fieldtrip toolbox (Oostenveld et al., 2011) and in-house-built scripts. Firstly, a low-pass filter at 40 Hz using a finite impulse response (FIR) filter with Kaiser window was applied to continuous MEG data. Then, the data were resampled to 200 Hz to save computational power and were epoched into 2-second segments to increase the signal-to-noise ratio. Around 1% of those 2-second segments were excluded as the corresponding auditory stimuli contained silent periods of more than 1 second. With the Fieldtrip automatic artifact rejection algorithm, we rejected trials with z-value higher than 100 before conducting independent component analysis (ICA). Using ICA, we identified and removed components corresponding to blinks, eye movements, cardiac activities, and residual noise from local train power. On average 7.6 ± 2.7 (SD) components were removed. The automatic artifact rejection algorithm was applied again to reject trials with z-value higher than 50.

For the further TRF (Section 2.3.3) and envelope reconstruction analyses (Section 2.3.5), another band-pass filter between 0.5 and 8 Hz was applied to the data (FIR filter with Kaiser window). For the coherence analysis, we re-segmented the preprocessed data into 4-sec segments to increase frequency resolution.

#### 2.3.3 Temporal response functions

The TRF analysis was done using the mTRF toolbox Version 2.0 (Crosse et al., 2016). TRFs were estimated by mapping speech features (e.g., envelopes, spectrograms) to neural responses of M/EEG. The mapping from stimulus to neural response is also known as “forward” modeling. The TRFs provide sensor-specific predictions and can indicate how a change of a stimulus feature (speech envelope) affects time-resolved neural activity at a certain location. In the context of a sensory system where the output is recorded by N recorded channels, we can assume that the instantaneous neural response *r*(*t, n*), which samples at times t = 1… T and at channel n, can be modeled over a convolution of the stimulus property, *s*(*t*), with a channel-specific TRF, *TRF*(*τ, n*). The TRF, *TRF*(*τ, n*), describes this transformation for a specified range of time lags, The response, *r*(*t, n*), can be modeled as: *r*(*t, n*) = ∑_*τ*_ *TRF*(*τ, n*)*s*(*t* − *τ*) + *ε*(*t, n*). The *ε*(*t, n*) is the residual response at each channel. Here, the TRF was estimated using ridge regression as follows, written in matrix format: *TRF* = (*S*^*T*^*S* + *λI*) ^−*1*^*S*^*T*^*r*. Where S is the lagged time series of the stimulus, *I* is the identity matrix, and *λ* is the regularization parameter (Crosse et al., 2016). The TRF was estimated over time lags ranging from -150 to 450 ms. In a 5-fold cross-validation procedure, trained TRFs are used to predict the left-out MEG response based on the corresponding speech envelope. The predicted response is then compared with the recorded neural data to assess MEG prediction accuracy. To optimize the model for predicting speech envelope, the values of the regularization parameter (*λ*) between 1 and 10^6^ was tuned using a leave-one-out cross-validation procedure. The *λ* value that produced the highest MEG prediction accuracy, averaged across trials and channels, was selected as the regularization parameter in each condition per participant. To increase signal to noise ratio, baseline normalization was applied using the time interval from -40 to 0 ms as baseline interval and computing the absolute change in TRF estimates with respect to the baseline interval. For further TRF sensor-level statistical analysis, we only used the gradiometers and calculated the combined planar gradient of the TRF estimates.

#### 2.3.4 Source Projection of Temporal Response Functions

To transform the TRF sensor data into source space, we used a template structural magnetic resonance image (MRI) from Montreal Neurological Institute (MNI) and warped it to the individual head shape (Polhemus points) to match the individual fiducials and head shape landmarks. A 3-dimensional grid covering the entire brain volume of each participant with a resolution of 1 cm was created based on the standard MNI template MRI. We then used a mask to keep only the voxels corresponding to the gray matter (1,457 voxels). The aligned brain volumes were further used to create realistic single-shell head models and lead field matrices (Nolte, 2003). By using the lead fields and a common covariance matrix (from all 2-second segments), common linearly constrained minimum variance (LCMV, Van Veen et al., 1997) beamformer spatial filter weights were computed based on an average covariance matrix estimated across all epochs. The regularization parameter was set to 20%. We then applied the spatial filter to the sensor-level TRF data.

#### 2.3.5 Speech envelope reconstruction

The speech envelope reconstruction was also done by using the mTRF toolbox (Crosse et al., 2016). We used a backward decoding model which uses a linear filter or decoder that optimally combines the MEG signals of different sensors in order to reconstruct the speech envelope. The decoder, *g*(*τ, n*), represents the linear mapping from the neural response, *r*(*t, n*), back to the stimulus, *s*(*t*). Reconstructed stimulus (speech envelope), *s*^(*t*) can be model as: *s*^(*t*) = ∑_*n*_ ∑_*τ*_ *r*(*t* + *τ, n*)*g*(*τ, n*). Analogous to the TRF approach, the decoder is computed as follows: *g* = (*R*^*T*^*R* + *λI*) ^−*1*^*R*^*T*^*s*, where *R* is the lagged time series of the response matrix, *r*.

The steps to obtain our measurement of neural speech tracking using this reconstruction method are the following. Firstly, the speech envelopes and MEG signals were down-sampled to 50 Hz to reduce the processing time. Secondly, we implemented a 5-fold cross-validation procedure and trained a linear decoder that combines the signals of all MEG channels and their time-shifted versions (integration window: -150-450 ms) on the MEG responses to the auditory stimuli. The decoder was then applied to the MEG responses to obtain reconstructed envelopes. The accuracy of reconstruction was measured by correlating the reconstructed envelope with the original envelope.

#### 2.3.6 Speech-brain phase coherence

We directly projected preprocessed sensor space data to source space using LCMV beamformer filters to obtain time series data of each brain voxel (Van Veen et al., 1997). The procedure of source projection is comparable to those for the TRF analysis in section 2.3.4. For calculating coherence between each brain voxel and the speech envelope, we then applied a frequency analysis to the 4-sec segments of all 6 conditions (original, 7-, 5-, 3-, 2-, 1-channel vocoded) using multi-taper frequency transformation (dpss taper: 1-25 Hz in 0.25 Hz steps, 4 Hz smoothing, no baseline correction).

In addition to canonical coherence analysis, we also looked into two sets of components of the coherence spectrum: periodic components (peak center frequency, peak bandwidth, peak height) and aperiodic components (aperiodic exponent, aperiodic offset) (Donoghue et al., 2020; Schmidt et al., 2021). These components were calculated using the FOOOF module with Python to compute aperiodic estimates and Gaussian model fits (Donoghue et al., 2020). The coherence data from 1 to 15 Hz was extracted from voxels in which the significant degradation effect was observed. The peak and aperiodic components of the coherence data were computed per voxel of each participant. The parameters used for modeling were from default setting (e.g., peak_width_limits: (0.5 12); max_n_peaks: inf; peak_threshold: 2.0). If R^2^ (between the input spectrum and the full model fit) of the residual model or error of the full model fit differed from the rest by more than 3 standard deviations, the result from the voxel was dropped. Peak and aperiodic components were then averaged across the rest of the voxels for each participant.

#### 2.3.7 Statistical analysis

For statistical comparisons, we first verified whether data distribution violated normality using the Shapiro-Wilk test (all *P*s > 0.5) and whether the data contained outliers with a 1.5 interquartile range criterion. We used parametric tests for data that obey normal distribution and outliers free; otherwise, non-parametric tests were used. If not mentioned specifically, multiple comparisons were corrected by using the false discovery rate method (FDR, Benjamini & Hochberg, 1995).

For the behavioral hit rates, the Friedman test was used to test the speech degradation effect across conditions. Paired Wilcoxon signed-rank tests were used to compare the hit rates between conditions. One-sample Wilcoxon signed-rank tests were used to test the hit rates against chance level (50%).

For the TRF sensor-level data, non-parametric cluster-based permutation tests (Maris & Oostenveld, 2007) implemented in the FieldTrip toolbox (Oostenveld et al., 2011) were used. The dependent sample F-statistic (“depsamplesFunivariate”) was used for cluster formation when analyzing the main effect of speech degradation with 0.05 alpha level. The cluster-level statistic via the Monte Carlo approximation with 0.05 alpha level was calculated as the maximum of the cluster-level summed F-values of each cluster with a minimum of 3 neighboring channels. The Monte-Carlo estimation was based on 10,000 random partitions, and the time window of interest was defined from 0 to 400 ms.

For the TRF source-level data, we used time windows from the clusters showing significant effects with the cluster-based permutation test in sensor-level analysis and ran the F-statistic where we averaged over the time window for each condition and compared the average across conditions. For contrast between conditions, we extracted TRF estimates from voxels with significant effect and ran paired Wilcoxon signed-rank tests.

For the stimulus reconstruction data, the correlation coefficient which represents reconstruction accuracy was Fisher z transformed for further statistical testing. Repeated-measure one-way ANOVA was used to test the degradation effect. Paired t-tests were used to compare the hit rates between conditions.

For the coherence data, the dependent sample F-statistic was used for cluster formation when analyzing the main effect of speech degradation with a 0.05 alpha level. The cluster - level statistic via the Monte Carlo approximation with 0.05 alpha level was calculated as the maximum of the cluster-level summed F-values of each cluster with a minimum of 3 neighboring channels. The Monte-Carlo estimation was based on 10,000 random partitions and the frequency window of interest was defined from 2 to 7 Hz. Paired Wilcoxon signed-rank tests were used for comparison between conditions.

For the above statistical tests, the corresponding effect size was calculated. For repeated-measure one-way ANOVA, partial eta square (η_p_^2^) was provided: η_p_^2^ = 0.01 indicates a small effect; η_p_^2^ = 0.06 indicates a medium effect; η_p_^2^ = 0.14 indicates a large effect. For paired t-tests, Cohen’s d (d) was provided: d = 0.2 indicates a small effect; d = 0.5 indicates a medium effect; d = 0.8 indicates a large effect. For the Friedman test, the Kendall’s W (W) was provided: W = 0.1 indicates a small effect; W = 0.3 indicates a medium effect; W = 0.5 indicates a large effect. For paired Wilcoxon signed-rank test and one-sample Wilcoxon signed-rank test, r was calculated as Z statistic divided by the square root of the sample size.: r = 0.1 indicates a small effect; r = 0.3 indicates a medium effect; r = 0.5 indicates a large effect.

To better understand functional characteristics of the TRF components, we calculated repeated measures correlations between TRF components and behavior hit rate, stimulus reconstruction accuracy as well as coherence results. The *rmcorr* function in the *rmcorr* package (Bakdash & Marusich, 2017) was used to calculate the correlation coefficient, 1,000-repetition bootstrapped 95% confidence interval, and p-value. Only the statistical tests using the Fieldtrip toolbox were conducted with MATLAB; the others were run with R software (version 3.6.2, R Development Core Team, 2019).

## 3. Results

### 3.1 Behavioral performance declines with decreased speech intelligibility

Sixteen of 24 healthy participants participated in both MEG and behavioral sessions, listening to audiobooks with 6-level of vocoded conditions (Figure 1; original, 7-, 5-, 3-, 2-, and 1-channel vocoded). At the end of each audio presentation, participants were required to choose which noun is the last noun they heard from the two nouns on the screen. We pooled the behavioral responses from the MEG session and the behavioral session (Figure 1C), which resulted in 12 behavioral responses in each condition. Participants’ task performance decreased as the intelligibility of the audio stimuli dropped. The mean hit rate was 100.0% ± 0% (SD) for the original stimuli, 88.5% ± 11.3% for the 7-ch vocoded, 83.3% ± 14.9% for the 5-ch vocoded, 65.6% ± 19.0% for the 3-ch vocoded, 58.9% ± 14.1% for the 2-ch vocoded, and 47.9% ± 11.6% for the 1-ch vocoded. A speech degradation effect was found across the six conditions [*𝒳*^2^(5) = 62.0, *p* = 4.69 × 10^−12^, Friedman test, Kendall’s W = 0.775]. Comparison within the six conditions showed that the hit rate for original stimuli was higher than all the other conditions (all *p*_*fdr*_ < 0.01, pairwise Wilcoxon signed-rank test, all effect size r > 0.80). The 7-ch vocoded condition had higher hit rates than 3-, 2-, and 1-ch conditions (all *p*_*fdr*_ < 0.01, all r > 0.25). The 5-ch condition had higher hit rates than 3-, 2-, and 1-ch conditions (all *p*_*fdr*_ < 0.01, all r > 0.60). The 3-ch condition had higher hit rates than the 1-ch condition (*p*_*fdr*_ = 0.01, r = 0.67). The 2-ch condition had higher hit rates than the 1-ch conditions (*p*_*fdr*_ = 0.034, r = 0.53). Except for the 1-ch vocoded condition (*p =* 0.38, r = 0.23, one-sample Wilcoxon signed-rank test), all the other conditions showed above-chance hit rates (all *p* < 0.03, all r > 0.56).

### 3.2 Temporal response functions show differential effects of vocoding on neural speech tracking

To investigate how a loss of spectral resolution influences speech tracking in a fine-grained temporal manner, we computed the temporal response functions (TRFs) between the speech envelope and the MEG signals in each condition from 24 participants.

The mean global field power in Figure 2A characterizes temporal evolution of TRF estimates in each condition. To investigate the effects of degraded speech on TRF, we ran a cluster-based permutation test with a time window of interest from 0 to 400 ms and observed a main effect of speech degradation in six clusters. These six clusters revealed bilateral effects in three time windows respectively (Figure 2A topographies): around 50-110 ms (termed as M50_TRF_, *p*_*cluster1*_ = 1.0 × 10^−4^, *p*_*cluster2*_ = 0.033), around 175-230 ms (termed as M200_TRF_, *p*_*cluster3*_ = 0.003, *p*_*cluster4*_ = 0.021), and around 315-380 ms (termed as M350_TRF_, *p*_*cluster5*_ = 0.001, *p*_*cluster6*_ = 0.035). We then projected the TRF sensor result to the source level (Figure 2B) and extracted TRF estimates from voxels showing degradation effect (Figure 2C). The statistical effect of M50_TRF_ on the source level was mainly around bilateral temporal gyrus, and the amplitude of the original condition showed the smallest effect compared to 7-ch (*p*_*fdr*_ *=* 2.21 × 10^−4^, pairwise Wilcoxon signed-rank test, effect size r = 0.77), 5-ch (*p*_*fdr*_ *=* 2.21 × 10^−4^, r = 0.79), 3-ch (*p*_*fdr*_ *=* 6.15 × 10^−4^, r = 0.71), 2-ch (*p*_*fdr*_ *=* 6.15 × 10^−4^, r = 0.71), and 1-ch (*p*_*fdr*_ *=* 2.21 × 10^−4^, r = 0.76).

**Figure 2.**
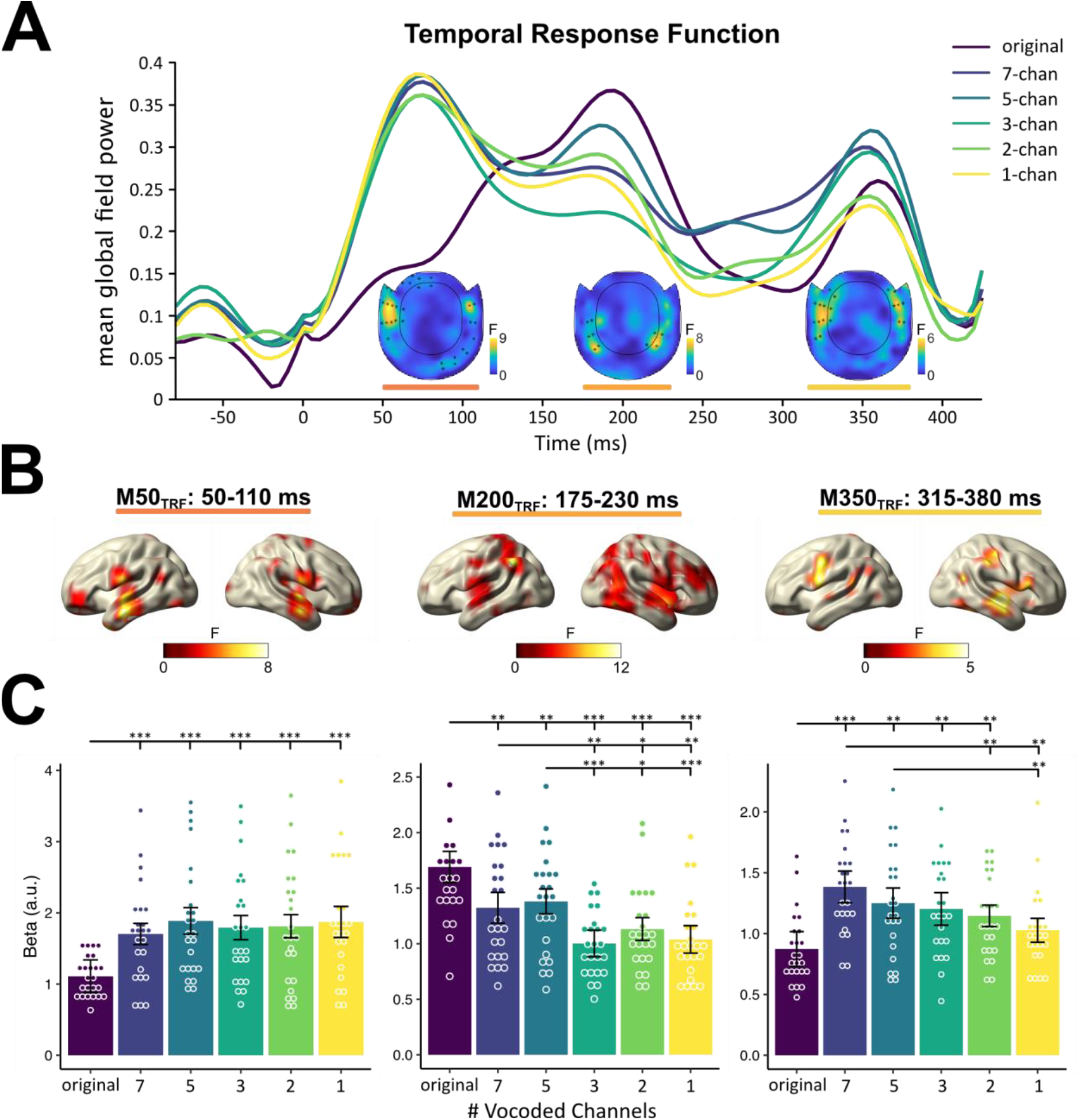
Temporal response functions (TRFs) of six-level degraded speech. (**A**) Global mean field of the TRF on the sensor level of each condition. The orange-yellow horizontal lines denote the time windows which show significant speech degradation effects. The topographies show the region of the speech degradation effect at each latency respectively. Sensors showing significant effects are denoted with asterisks. (**B**) Source localizations of the degradation effects for each time window. (**C**) Individual TRF estimates of the six conditions extracted at voxels showing significant degradation effect. Bars represent 95% confidence intervals. *p*_*fdr*_ < .05*, *p*_*fdr*_ < .01**, *p*_*fdr*_ < .001***

The statistical effect of M200_TRF_ on the source level was around the bilateral parietal and temporal region. The source estimates overall attenuated as the intelligibility decreased. The original condition showed higher amplitude than 7-ch (*p*_*fdr*_ *=* 0.003, r = 0.62), 5-ch (*p*_*fdr*_ *=* 0.002, r = 0.65), 3-ch (*p*_*fdr*_ *=* 1.78 × 10^−6^, r = 0.88), 2-ch (*p*_*fdr*_ *=* 4.47 × 10^−6^, r = 0.86), and 1-ch (*p*_*fdr*_ *=* 1.96 × 10^−5^, r = 0.82). The 7-ch condition had higher amplitude than 3-ch (*p*_*fdr*_ *=* 0.003, r = 0.62), 2-ch (*p*_*fdr*_ *=* 0.026,, r = 0.47) and 1-ch (*p*_*fdr*_ *=* 0.006, r = 0.57). The 5-ch condition had higher amplitude than 3-ch (*p*_*fdr*_ *=* 8.34 × 10^−4^, r = 0.69), 2-ch (*p*_*fdr*_ *=* 0.011, r = 0.54), and 1-ch (*p*_*fdr*_ *=* 7.69 × 10^−4^, r = 0.71).

The strongest responses of source of M350_TRF_ were observed in the vocoded but still comprehensible conditions (i.e., 7-ch, 5-ch, and 3-ch conditions); the weakest response showed in the original condition. The 7-ch condition had a higher amplitude than 2-ch (*p*_*fdr*_ *=* 0.009, r = 0.58), 1-ch (*p*_*fdr*_ *=* 0.002, r = 0.79), and the original condition (*p*_*fdr*_ *=* 3.70 × 10^−4^, r = 0.78). The 5-ch condition had a higher amplitude than the 1-ch (*p*_*fdr*_ *=* 0.009, r = 0.57) and original condition (*p*_*fdr*_ *=* 0.009, r = 0.60). The 3-ch and 2-ch conditions had a higher amplitude than the original condition (*p*_*fdr*_ *=* 0.009 and 0.009, r = 0.61 and 0.57, respectively).

Overall, speech intelligibility did not modulate the temporal response function in the same manner over time. Modulation was observed at three intervals, and only the effect around the middle interval (M200_TRF_) was related to speech intelligibility in a relatively straight-forward manner, namely that only the amplitude of M200_TRF_ decreased with the reduced intelligibility. These differential TRF patterns could be attributed to either general speech feature gain modulation or tracking in specific speech features e.g. amplitudes of speech envelope or syllables. In order to better understand these complex temporal patterns, we quantified other neural tracking measures and related them to the main TRF effects, which is described in the following sections.

### 3.3 Accuracy of the speech envelope reconstruction generally declines with decreased speech intelligibility

The envelope reconstruction accuracy was evaluated by correlation coefficient as reconstruction accuracy between the original envelope and the reconstructed envelope (Figure 3A). A speech degradation effect on reconstruction accuracy was observed [repeated measure one-way ANOVA, F(5,115) =9.16, *p* = 2.34 × 10^−7^, η_p_^2^ = 0.29]. The original condition had higher accuracy than the 2-ch [t(23) = 3.30, *p*_*fdr*_ = 0.007, Cohen’s d = 0.67] and 1-ch conditions [t(23) = 4.87, *p*_*fdr*_ = 4.88 × 10^−4^, d = 0.99]. The 7-ch condition had higher accuracy than the 2-ch [t(23) = 3.48, *p*_*fdr*_ = 0.005, d = 0.71] and 1-ch conditions [t(23) = 3.82, *p*_*fdr*_ = 0.003, d = 0.78]. The 5-ch condition had higher accuracy than the 3-ch [t(23) = 3.58, *p*_*fdr*_ = 0.005, d = 0.73], 2-ch [t(23) = 4.47, *p*_*fdr*_ = 8.65 × 10^−4^, d = 0.92], and 1-ch conditions [t(23) = 5.39, *p*_*fdr*_ = 2.68 × 10^−4^, d = 1.10]. In sum, the reconstruction accuracy overall declined as the speech intelligibility decreased, while no differences were found among the original, 7-ch, and 5-ch conditions.

**Figure 3.**
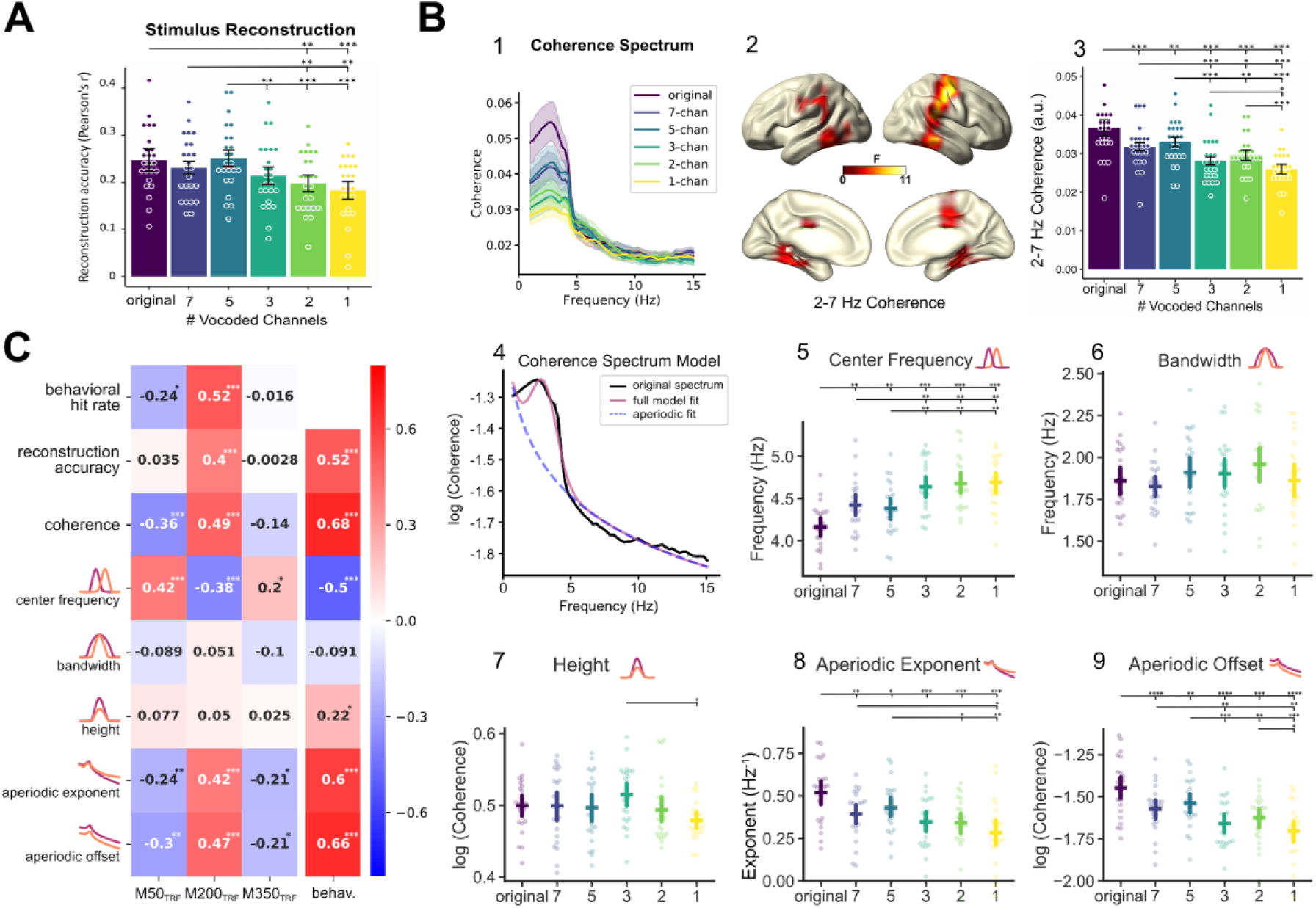
Neural tracking of speech intelligibility measured via stimulus reconstruction and coherence. (**A**) Accuracy (Pearson’s correlation coefficient) of stimulus reconstruction for each condition. (**B**) Coherence and components of the coherence spectrum in six conditions. **1**, Frequency spectrum of the coherence for the six conditions averaged across all voxels. **2**, Source localization of degradation effect on coherence across six conditions in bilateral temporal, inferior frontal, and parietal regions. **3**, Individual coherence values of the six conditions extracted from voxels showing significant effects. **4**, Example of model fitting for components of one coherence spectrum. **5-9**, Estimated peak center frequency, peak bandwidth, peak height, aperiodic exponent, and aperiodic offset of coherence spectrum in six conditions. Bars represent 95% confidence intervals. (**C**) Correlations between the TRF components and the behavioral hit rate, reconstruction accuracy, coherence as well as components of coherence spectrum show that M200_TRF_ moderately correlated with hit rate, reconstruction accuracy, coherence, center frequency, exponent and offset which suggest that M200TRF can be another neural index of speech intelligibility. *p*_*fdr*_ < .05*, *p*_*fdr*_ < .01**, *p*_*fdr*_ < .001***

### 3.4 Speech-brain coherence declines with decreased speech intelligibility

Speech-brain coherence is likely the most common measure to quantify neural tracking. A degradation effect of speech was found around bilateral temporal, inferior parietal and inferior frontal regions from the broadband 2-7 Hz coherence analysis (Figure 3B-2; cluster-corrected dependent-sample F-test; *p*_*cluster1*_ = 9.99 × 10^−4^, *p*_*cluster2*_ = 0.018). Within these areas, the strongest effect was found in the original condition, and no differences were found between 7-ch and 5-ch as well as between 3-ch and 2-ch condition (Figure 3B-3). The original condition showed stronger coherence than the 7-ch (*p*_*fdr*_ *=* 1.27 × 10^−4^, pairwise Wilcoxon signed-rank test, effect size r = 0.74), 5-ch (*p*_*fdr*_ *=* 0.005, r = 0.58), 3-ch (*p*_*fdr*_ *=* 2.23 × 10^−6^, r = 0.86), 2-ch (*p*_*fdr*_ *=* 4.08 × 10^−5^, r = 0.79) and 1-ch condition (*p*_*fdr*_ *=* 8.92 × 10^−7^, r = 0.88). The 7-ch condition showed stronger coherence than the 3-ch (*p*_*fdr*_ *=* 6.86 × 10^−5^, r = 0.76), 2-ch (*p*_*fdr*_ *=* 0.02,, r = 0.48) and 1-ch conditions (*p*_*fdr*_ *=* 8.92 × 10^−7^, r = 0.88). The 5-ch condition showed stronger coherence than the 3-ch (*p*_*fdr*_ *=* 8.94 × 10^−6^, r = 0.83), 2-ch (*p*_*fdr*_ *=* 0.003, r = 0.61), and 1-ch conditions (*p*_*fdr*_ *=* 1.19 × 10^−6^, r = 0.87). The 3-ch condition showed stronger coherence than the 1-ch (*p*_*fdr*_ *=* 0.039, r = 0.43). The 2-ch condition showed stronger coherence than the 1-ch (*p*_*fdr*_ *=* 4.31 × 10^−5^, r = 0.78). In sum, speech-brain coherence overall declined as the speech intelligibility decreased, while no differences were found between the 7-ch and 5-ch as well as between the 3-ch and 2-ch condition.

### 3.5 The peak center frequency, aperiodic exponent, and aperiodic offset are modulated by speech intelligibility

To complement our finding in the TRFs, we further extracted periodic components (peak center frequency, peak bandwidth, peak height) and aperiodic components (exponent, offset) from the broadband coherence spectrum.

We extracted the coherence data from 1 to 15 Hz from the voxels in which the significant degradation effect was observed from 2 to 7 Hz in the previous section. The peak and aperiodic components were computed per voxel of each participant (Figure 3B-4). The peak and aperiodic components were then averaged across voxels which showed a significant degradation effect for each participant. Interestingly, we observed that the peak center frequency accelerated (mean center frequency from 4.17 Hz to 4.69 Hz) for less intelligible speech [*𝒳*^2^(5) = 41.0, *p* = 9.49 × 10^−8^, Friedman test, Kendall’s W = 0.78] (Figure 3B-5). This finding thereby fits with the idea proposed by Schmidt et al (2021) that the speech tracking shifts from the more linguistic level (syllabic rate of the speech 4.3 Hz) to the more acoustic level (envelope modulation rate ∼5 Hz). The aperiodic components (exponent & offset) decreased as the speech intelligibility decreased [exponent: *𝒳*^2^(5) = 35.2, *p* = 1.39 × 10^−6^, W = 0.29; offset : *𝒳*^2^(5) = 53.2, *p* = 3.07 × 10^−10^, W = 0.44] (Figure 3B-8, 3B-9). A borderline degradation effect was observed in the peak height [*𝒳*^2^(5) = 11.2, *p* = 0.048, W = 0.09] (Figure 3B-7), and no degradation effect was found in the peak bandwidth [*𝒳*^2^(5) = 6.43, *p* = 0.26, W = 0.05] (Figure 3B-6). In sum, our results show that with stronger vocoding, the peak center frequency shifted toward the acoustic level and the aperiodic components decreased.

### 3.6 The TRF finding highly correlates with behavioral and other neural measures of neural speech tracking

To relate our findings in TRF components with the other measures, we computed the repeated measures correlation coefficients between TRF components, behavior measurement, and the other neural indices (stimulus reconstruction accuracy, coherence, and components of coherence spectrum). The repeated measures correlation coefficients between the TRF components and the other measures are shown in Figure 3C.

While the higher M50_TRF_ was mildly correlated with the lower behavioral hit rates [*r*_rm_(79) = - 0.24 (−0.41 to -0.05), *p* = 0.033], the enhancement in M50_TRF_ was correlated with the center frequency shifting to the acoustic level [*r*_rm_(119) = 0.42 (0.25 to 0.56), *p* = 1.85 × 10^−6^]. Besides, the higher M50_TRF_ was also mildly correlated with the lower coherence [*r*_rm_(119) = -0.36 (−0.52 to -0.19), *p* = 6.34 × 10^−5^], and the lower aperiodic components [exponent: *r*_rm_(119) = -0.24 (−0.37 to -0.07), *p* = 0.008; offset: *r*_rm_(119) = -0.30 (−0.44 to -0.12), *p* = 0.001].

Of the three TRF components, M200_TRF_ can best predict behavioral performance [*r*_rm_(79) = 0.52 (95% CI: 0.40 to 0.65), *p* = 5.25 × 10^−7^]. Furthermore, the attenuation in M200_TRF_ was correlated with the center frequency shifting from the linguistic to the acoustic level [*r*_rm_(119) = -0.38 (−0.50 to -0.24), *p* = 1.74 × 10^−5^], which was opposite to M50_TRF_. In addition, M200_TRF_ was also positively correlated with the reconstruction accuracy [*r*_rm_(119) = 0.40 (0.29 to 0.52), *p* = 4.42 × 10^−6^], the coherence [*r*_rm_(119) = 0.49 (0.35 to 0.61), *p* = 1.21 × 10^−8^], and the aperiodic components [exponent: *r*_rm_(119) = 0.42 (0.26 to 0.54, *p* = 2.09 × 10^−6^; offset: *r*_rm_(119) = 0.47 (0.33 to 0.59), *p* = 4.88 × 10^−8^].

Compared to M50_TRF_ and M200_TRF_, the nonlinear pattern of M350_TRF_ showed mild correlation to only three neural measures, which was correlated positively with the center frequency [*r*_rm_(119) = 0.20 (0.03 to 0.36), *p* = 0.030] and negatively with the aperiodic components [exponent: *r*_rm_(119) = -0.21 (−0.34 to -0.07), *p* = 0.019; offset: *r*_rm_(119) = -0.21 (−0.35 to -0.06), *p* = 0.022].

We also related the behavior results to the other neural measures: The better behavior hit rates were highly correlated with the higher envelope reconstruction accuracy [*r*_rm_(79) = 0.52, 95% CI: 0.35 to 0.67, *p* = 8.70 × 10^−7^], the higher coherence [*r*_rm_(79) = 0.68 (0.54 to 0.76), *p* = 4.52 × 10^−12^], the higher aperiodic components [exponent: *r*_rm_(79) = 0.60 (0.46 to 0.71), *p* = 2.99 × 10^−9^; offset: *r*_rm_(79) = 0.66 (0.56 to 0.77), *p* = 1.64 × 10^−11^] but with the lower center frequency [*r*_rm_(79) = -0.50 (−0.65 to -0.34), p = 2.16 × 10^−6^]. The correlation with the peak height was weak but statistically significant [*r*_rm_(79) = 0.22 (0.03 to 0.42), p = 0.046].

Together, among the three TRF components, only M200 was highly correlated with the behavioral results, and both of them were correlated with reconstruction accuracy, speech-brain coherence, peak center frequency, and aperiodic components. Interestingly, both M50 and M200 were correlated with the shifting center frequency but in the opposite direction. These findings depicted how vocoded speech affects neural speech tracking over time.

## 4. Discussion

Neural speech tracking is modulated by the spectral details of speech. However, the reported pattern of the intelligibility modulation has been mostly computed in a “static” manner [e.g. stimulus reconstruction (Verschueren et al., 2019) and speech-brain coherence (Hauswald et al., 2020)]. Here, we aimed to explore how the human brain represents different levels of degraded speech in a time-resolved manner. For this reason, we used original (i.e. clear and unaltered) speech and 5 levels of vocoded speech as stimuli and computed time-resolved temporal response functions (TRFs). Secondly, we related TRF findings with behavioral and other neural measures (stimulus reconstruction, speech-brain coherence, and components of broadband coherence spectra) of neural speech tracking. Overall, the behavioral performance declined with decreased speech intelligibility as expected. Our results showed that various levels of degraded speech differentially modulate the TRF around three time-windows: 50-110 ms (M50_TRF_), 175-230 ms (M200_TRF_), and 315-380 ms (M350_TRF_). Interestingly, both M50 and M200 are correlated with the shifting center frequency of the coherence spectrum, however, in the opposite direction. These findings suggest that the distinctive effects on TRF are linked to altered tracking of speech features.

### 4.1 Neural tracking and speech intelligibility

In our neural measures, the modulation of the M200_TRF_, the speech brain coherence and the aperiodic components of the broadband coherence spectra support the notion that the neural speech tracking declines as the speech intelligibility decreases.

The M200_TRF_ is the TRF component that reflects the behavioral response best, i.e. captures the speech intelligibility (for the discussion of M50_TRF_ and M350_TRF_, see section 4.2). Its amplitude decreases as the speech intelligibility declines (Figure 2C-central). This relationship is also shown by the similarity of the correlational patterns of the M200_TRF_ and the behavioral performance with the other measures (see Figure 3C middle and right column). This finding of pointing to the relevance of speech intelligibility (measured behaviorally) of the M200 _TRF_ is consistent with other studies, e.g. in an EEG study with multi-talker and vocoded speech design, greater TRF deflection around 200 ms was also observed in an attended-nonvocoded condition compared to an attended-vocoded condition (Kraus et al., 2021). Using conventional ERP analysis, greater P200 amplitude has been also observed in clear speech compared to degraded speech (Strauß et al., 2013) and in words pronounced with native speech than words pronounced with foreign-accented speech (Romero-Rivas et al., 2015). Besides, the P200 amplitude has been positively correlated with the successful extraction of phonetic/phonological information from vowels or words (De Diego Balaguer et al., 2007; Reinke et al., 2003). All the above results indicate that the higher amplitude of this component is, the higher speech intelligibility is. Furthermore, the M200_TRF_ effect is located mainly around inferior parietal cortex, superior temporal gyrus, and inferior frontal cortex, consistent with prior fMRI work on the effect of speech intelligibility (Obleser & Kotz, 2010).

The association between the M200_TRF_ and the peak center frequency of the coherence spectrum also supports the notion that M200_TRF_ can be an index of speech intelligibility (Schmidt et al., 2021). The shifting center frequency reflects the neural speech tracking from the (linguistic-level) syllabic rate of the speech (4.3 Hz) to the (acoustic-level) envelope modulation rate (∼5 Hz). This is not merely due to the physical characteristics of the acoustic stimulus as the reduction of spectral details (from clear to 1-ch vocoded) in the audio stimuli results in the decrease of peak frequency in the amplitude modulation of the stimuli (from 5.61 to 4.83 Hz). However, the center frequency of the speech brain coherence, on the other hand, increased (from 4.17 to 4.69 Hz). This phenomenon also highlights that the brain is not exclusively driven by the rhythm of external stimuli.

The speech-brain coherence was also highly correlated with the behavior performance, which reflects speech intelligibility, in line with Schmidt et al. (2021). However, the coherence result was different from Hauswald et al. (2020), from which we re-analyzed the data. In Hauswald et al. (2020), the coherence result showed an inverted U-shaped pattern: The highest coherence showed in the marginally intelligible (5-ch vocoded) condition. After re-analyzing the data, we found that the difference was attributed to distinct high-pass filter settings (online 0.1 Hz here instead of 1 Hz). This could also explain the finding of lower inter-trial phase correlation in delta band in the original and 8-ch vocoded conditions compared to the 4-ch vocoded condition in Ding et al. (2014) as the high-pass filtered was also set at 1 Hz.

A speech degradation effect was found in the aperiodic components (exponent & offset), consistent with Schmidt et al. (2021). Here, we demonstrate that the aperiodic components were highly correlated with the behavioral performance, which suggests that the aperiodic components can be reliable biomarkers for speech processing. Overall, we again highlight the importance of both the periodic and aperiodic components of the coherence spectra as they can provide more detailed information of speech processing than the coherence alone. As the periodic and aperiodic components of the frequency power spectrum have been shown to be biological markers in aging (Dave et al., 2018; Voytek et al., 2015) and mental disorders (Molina et al., 2020; Ostlund et al., 2021; Robertson et al., 2019), it is possible that this can also be the case for components of the coherence spectrum. Future studies could test whether those components could offer more insight into more clinically-relevant participants, e.g. in individuals with hearing impairment.

### 4.2 Neural speech tracking could be affected by other potential factors

The modulation of spectral details in speech signals not only affects speech intelligibility but also influences e.g. attention load, memory load, and perceptual learning (Mattys et al., 2012). Our findings in M50_TRF_ and M350_TRF_ also argue that speech intelligibility can be modulated by other factors.

M50_TRF_, the earliest effect occurred around 50 ms and clearly differentiated all vocoded conditions from the non-vocoded one (Figure 2C-left). The finding of this early component is in line with previous studies showing that the TRF as well as ERP amplitudes of this early peak were greater in vocoded conditions compared to intact conditions (Ding et al., 2014; Kraus et al., 2021; Strauß et al., 2013). Besides, a comparable effect was also found for louder conditions when compared to soft-spoken conditions (Verschueren et al., 2021). While this M50_TRF_ peak thus seems to follow physical manipulations of the speech signal, Ding & Simon (2012) could not find attentional modulations earlier than ∼100 ms. Taken together, we propose that this early effect might be modulated by sensory gain on acoustic properties of the auditory stimuli rather than task difficulty as there is marked distinction between clear (normal) and spectral-degraded (abnormal) speech. In addition, our source result showing the prominent effect around bilateral primary auditory cortices further support the notion that the speech processing is rather early at this stage (de Heer et al., 2017; Khalighinejad et al., 2021). It is worth noting that a similar effect on this early component was also observed for age both in pure tones (Herrmann et al., 2022) and in speech (Brodbeck et al., 2018), with the older participants showing an increased amplitude compared to the younger ones. The potential neural mechanism in these cases can be the altered balance between inhibitory and excitatory neural mechanisms in the cortex during aging. Furthermore, we also found that the higher M50_TRF_ is associated with the increased center frequency of coherence as the speech intelligibility decreased. This finding highlights that M50_TRF_, which reflects the distinctions of acoustic information between original and vocoded speech, can also play a role in shifting center frequency for degraded speech.

The effect of speech degradation on TRF around 350 ms (M350_TRF_) shows an inverted U-shaped pattern across six conditions: the maximal amplitude found in 7-ch condition, followed by 5-ch, 3-ch, 2-ch, 1-ch, and original (Figure 2C-right). The M350_TRF_ effect was source-localized mainly around the left inferior frontal cortex. This M350_TRF_ shares temporal and source characteristics with the N400 which is seen as a potential marker of semantic integration (for review, see Kutas & Federmeier, 2011; Lau et al., 2008). It is possible that the strongest effect observed in the 7-ch vocoded condition resulted from that the speech was still understandable and predictable while the acoustic input was violated. The effect reduced in the clear speech as the acoustic input was as predicted. Furthermore, the effect declined in the more degraded speech as the speech became less comprehensible and less predictable. Accordingly, a similar nonlinear N400 effect was also found in a previous study manipulating 6 levels of signal to noise ratio (Jamison et al., 2016). Furthermore, lexical-semantic processing is modulated when the spectral properties of the speech are disrupted as shown by Obleser & Kotz (2011) and Strauß et al. (2013). Taken together, previous research and our results seem to indicate that the M350_TRF_ is an index of late-stage speech processing (e.g. semantic integration), whereas more research directly addressing this question is needed.

Based on our time-resolved findings in TRF components, we propose that envelope reconstruction is not a robust measure for the modulation of speech intelligibility on neural speech tracking as the result of envelope reconstruction is rather static. This argument is supported by results from ours and previous studies (Decruy et al., 2020; Presacco et al., 2019). In our results, no differences were observed among the conditions that can be understood very easily to mildly challenging (original, 7-ch, and 5-ch conditions). In Decruy et al. (2020) and Presacco et al. (2019), higher reconstruction accuracy did not fully reflect better speech understanding especially when comparing hearing impaired populations to healthy control groups.

### 4.3 Conclusions

In this study, we demonstrate that at least three neural processing stages (M50_TRF_, M200_TRF_, and M350_TRF_) are affected when processing continuous degraded speech. Only M200_TRF_ highly correlates with behavior and other neural measures (i.e., speech brain coherence) of speech intelligibility. We also indicate that both M50_TRF_ and M200_TRF_ can reflect shifts in the neural speech tracking from more linguist level to more acoustic level as speech intelligibility declined which is supported by the effect of center frequency of coherence. In general, using the temporal response function combined with parametrization of coherence spectra, we demonstrate the temporal dynamics of neural speech tracking and provide potential explanations for the inconsistent literature.

## Acknowledgements

We thank the participants in this study as well as Manfred Seifter and Bernadette Eckart for their support during data collection.

## Data/Code Availability

The data and code necessary for generating the figures and computing statistics will be shared in the corresponding author’s gitlab repository: https://gitlab.com/yapingchen. Access to raw data will be made available upon reasonable request to the corresponding author.

## Author contributions

Funding acquisition: NW, AK, AH, SR

Conceptualization: NW, FS, YC, AH, AK

Data curation: YC, FS, AH

Formal Analysis: YC, FS, AH

Supervision: NW, AH, AK

Visualization: YC, FS

Writing—original draft: YC

Writing—review & editing: AH, FS, AK, NW

